# Conservation of sex-linked markers among conspecific populations of a viviparous skink, *Niveoscincus ocellatus,* exhibiting genetic and temperature dependent sex determination

**DOI:** 10.1101/257428

**Authors:** Peta Hill, Christopher P. Burridge, Tariq Ezaz, Erik Wapstra

## Abstract

Sex determination systems are exceptionally diverse and have undergone multiple and independent evolutionary transitions among species, particularly reptiles. However, the mechanisms underlying these transitions have not been established. Here we tested for differences in sex-linked markers in the only known reptile that is polymorphic for sex determination system, the spotted snow skink, *Niveoscincus ocellatus,* to quantify the genomic differences that have accompanied this transition. In a highland population, sex is determined genetically, whilst in a lowland population, offspring sex ratio is influenced by temperature. We found a similar number of sex-linked loci in each population, including shared loci, with genotypes consistent with male heterogamety (XY). However, population-specific linkage disequilibrium suggests greater divergence of sex chromosomes in the highland population. Our results suggest that transitions between sex determination systems (GSD and TSD-like systems) can be facilitated by subtle genetic differences.

## Introduction

Sex determination controls whether the embryonic gonads develop into testes or ovaries. Central to the development of sexually reproducing organisms, sex determination should be under strong selection with highly conserved processes and limited evolutionary lability (Uller, et al. 2007). Contrary to these expectations, systems of sex determination are surprisingly diverse, and therefore there is substantial interest in their evolution (Capel 2017; Ezaz, et al. 2006; O’Meally, et al. 2012). Vertebrate sex can be determined by genes (genetic sex determination; GSD), the environment (environmental sex determination; ESD), or via interactions between the two (Sarre, et al. 2004; Shine, et al. 2002; Valenzuela, et al. 2003). Furthermore, an extraordinary number of evolutionary transitions between these modes have occurred unpredictably across vertebrate evolution (Bachtrog, et al. 2014; Janzen, et al. 2006; Pokorna, et al. 2016). Sex determination also directs population sex ratio, an important demographic parameter that has implications for population persistence (Boyle, et al. 2014).

The mechanisms underlying sex determination systems are diverse. While a master genetic “switch” directs gonadogenesis in GSD species, it can manifest as a single nucleotide polymorphism (SNP; e.g. Takifugu rubripes; Kamiya, et al. 2012), a dominant single gene system (e.g. Mammalian *SRY;* Koopman, et al. 1990), a single gene dosage system (e.g. Avian *DMRT1*; Smith, et al. 2004), or methylation status of genes or their promoters (e.g. half-smooth tongue sole; Chen, et al. 2014). GSD is ubiquitous in endotherms and amphibians, and is found throughout lineages of reptiles and fish (Ezaz, et al. 2006; Quinn, et al. 2011; Sarre, et al. 2011). Environmental control of sex occurs in many ectotherms (Adkins-Regan, et al. 2014; Bull 1980; Ewert, et al. 2005), with temperature determining sex in many reptiles (temperature dependent sex determination, TSD). The environment can also act to over-ride the genetic influence of sex determination in a predominantly GSD species (GSD plus environmental effects, GSD+EE: Valenzuela, et al. 2003) with a temperature over-ride described in reptiles (Holleley, et al. 2015; Radder, et al. 2008; Shine, et al. 2002). Many reptiles may possess an environmental over-ride rather than strict GSD because sex determination is a continuous rather than dichotomous trait (Sarre, et al. 2004).

In pure TSD taxa sex ratios are close to all male or all female at sex-specific developmental temperatures, whilst at a very narrow pivotal temperature range they can be a mix of male and female (Ewert, et al. 2005; Lang, et al. 1994). A temperature over-ride of sex determination presents as sex ratios at 50:50 across a broad pivotal temperature range with deviations occurring outside this range (Holleley, et al. 2015; Shine, et al. 2002). Uncovering sex-linked genetic sequence in a species previously shown to have TSD places this species on the continuum between GSD and TSD (Sarre, et al. 2004; Valenzuela, et al. 2014). TSD has been extensively studied in oviparous reptiles (Georges 1989; Harlow, et al. 2000; Lang, et al. 1994; Valenzuela, et al. 2014), where offspring sex is labile until after the middle third of embryonic development (Shine, et al. 2007), and is mediated by nest temperature. Viviparity was traditionally considered incompatible with any form of temperature influence on sex determination (Bull 1980), yet it has recently been described in a handful of reptiles (Robert, et al. 2001; Wapstra, et al. 2004; Zhang, et al. 2010), with the temperature signal mediated by maternal basking behaviour.

The correlation between sex determination system and the presence or absence of differentiated sex chromosomes—chromosomes that differ morphologically between males and females—is surprisingly weak (Sarre, et al. 2004; Vicoso, et al. 2013b). When present, vertebrate sex chromosomes are remarkably diverse, even between closely related taxa (Bachtrog, et al. 2014; Ezaz, et al. 2017; Georges, et al. 2010), suggesting that contemporary sex chromosomes have multiple evolutionary origins (Ezaz, et al. 2009a; Kawai, et al. 2009; Matsubara, et al. 2006). Heterogamety for sex chromosomes can occur in males (XY, e.g., mammals) or females (ZW, e.g., birds), and sex chromosomes can be hetero- or homomorphic, regardless of sex determination mechanism (GSD, TSD or GSD+EE). Understanding how these multiple evolutionary transitions in sex determination have occurred requires exposing the mechanisms that underpin them at a molecular level; the degree to which sex chromosomes participate in, or are a product of, transitions between sex determination systems remains a key knowledge gap.

Dosage models have been used to explain both environmental influence on sex, and transitions in sex determination systems and sex chromosomes (Ezaz, et al. 2009b; Quinn, et al. 2007; Quinn, et al. 2011). Under a dosage model, one sex is determined when the product of a homogametic genotype reaches a certain threshold. If the gene product that determines sex possesses thermal sensitivity, it is possible for a heterogametic genotype to reach the same threshold, or a homogametic genotype to not, resulting in the reversal of genotypic sex and the bias of sex ratio towards the sex most likely to benefit from the environment experienced (Charnov, et al. 1977). Dosage models can also explain transitions in sex determination; selection on the threshold for sex can result in transitions between GSD and TSD, and between ZW and XY heterogamety if sex determination acquires temperature sensitivity (Quinn, et al. 2011). A transition in heterogamety can also occur via the invasion of a novel sex determining locus when existing sex chromosomes are undifferentiated (Bachtrog, et al. 2014; Schartl 2004). Sex chromosomes can also be lost during transitions from GSD to TSD (Holleley, et al. 2015).

Reptiles exhibit high diversity in sex determination systems and sex chromosome morphology and homology (Ezaz, et al. 2009b; Giovannotti, et al. 2010; Matsubara, et al. 2014; Norris 2003; Shine, et al. 2002), and therefore represent a valuable group for the study of transitions in sex determination and sex chromosome systems. However, incipient transition in sex determination has been documented only within one reptile, the viviparous spotted snow skink *Niveoscincus ocellatus* (Cunningham, et al. 2017; Pen, et al. 2010), representing a powerful study system. A highland population has GSD, whilst in a lowland population, temperature influences the sex of offspring in a TSD-like pattern of sex ratio bias These populations diverged recently, within the last million years (Cliff, et al. 2015). Divergent natural selection on sex determination caused by climatic effects on lizard life history and variation in the size of inter-annual temperature fluctuations is driving this transition (Pen, et al. 2010). Warmer years result in early birth in both populations but sex ratios respond to temperature only in the lowland (Cunningham, et al. 2017). Sex ratios in the lowland are female biased in warm years and male biased in cold years. Lowland females, but not males derive a selective advantage from being born early because birth date influences the onset of maturity and this is important for females (Wapstra, et al. 2004). In the highland sex ratios do not vary from parity regardless of temperature because birth date does not predict the onset of maturity in this population (Pen, et al. 2010). In addition, higher inter-annual variation in climate in the highland is thought to favour GSD because it prevents extreme sex ratios (Pen, et al. 2010). This establishes an adaptive explanation for intra-specific divergence in sex determination systems. However, knowledge gaps exist surrounding the mechanism of this transition and the background with respect to sex chromosome evolution. Modelling suggests divergence among populations in genes that control sex determination: loss of function in the lowland population, attainment of function in the highland population, or a combination of both (Pen, et al. 2010).

The aim of this study was to quantify the divergence of genomic regions associated with sex (sex-linked markers) in populations of *N. ocellatus* that have recently diverged in sex determination system. Explicitly, we test whether the two populations differ in the numbers of sex-linked markers and the levels of linkage disequilibrium around them. We discuss our findings with regard to sex determination and sex chromosome evolution.

## Materials and Methods

### Study species

*Niveoscincus ocellatus* is a small (60 to 80 mm snout-vent length, 3–10g) viviparous skink endemic to Tasmania, with a broad altitudinal distribution from sea level to 1200m (Wapstra, et al. 1999). Two study populations represent the climatic extremes of this species’ range: a cool temperate lowland population (42 34’S, 147 52’ E; elevation 50m; hereafter ‘lowland population’) and a cold temperate, sub-alpine population (41 51’S, 146 34’E; elevation 1200m; hereafter ‘highland population’). Reproduction follows a similar pattern in both populations; females reproduce annually, and the reproductive cycle is completed in one season (Wapstra, et al. 1999). Gestation occurs in spring and parturition in summer. Long term data on these populations consistently documents their divergent sex determination systems (Cunningham, et al. 2017; Wapstra, et al. 2004; Wapstra, et al. 2001).

### Genotyping by sequencing

Approximately 2–4mm of tail tip was sampled from 44 highland individuals (23 males,

21 females) and 44 lowland individuals (24 males and 20 females) during the 2014–15 season. Males were sexed in the field by hemipene eversion, and all females were observed to later give birth. DNA extractions and sequencing were performed using DArTseq^™^ (Diversity Arrays Technology PTY, LTD), a high-throughput genotyping by sequencing method (Kilian, et al. 2012) that employs genomic complexity reduction using restriction enzyme pairs. This technology successfully developed a series of sex-linked markers in the frog *Rana clamitans* (Lambert, et al. 2016). DNA was digested using *PstI* and *SphI.* Ligation reactions were then performed using two adaptors: a *PstI* compatible adaptor consisting of Illumina flow-cell attachment sequence, sequencing primer sequence and a unique barcode sequence, and a *SphI* compatible adaptor consisting of an Illumina flow-cell attachment region. Ligated fragments were then PCR amplified using an initial denaturation at 94°C for 1 min, followed by 30 cycles of 94°C for 20 sec, 58°C for 30 sec, and 72°C for 45 sec, with a final extension step at 72°C for 7 min. Equimolar amounts of amplification products from each individual were pooled and subjected to Illumina’s proprietary cBot (http://www.illumina.com/products/cbot.html) bridge PCR followed by sequencing on an Illumina Hiseq2000. Single read sequencing was run for 77 cycles.

Sequences were processed using proprietary DArTseq analytical pipelines (Ren et al. (2015). Initially, the Hiseq2000 output (FASTQ file) was processed to filter poor quality sequences. Two different thresholds of quality were applied. For the barcode region (allowing parsing of sequences into specific sample libraries), we applied more stringent selection (minimum phred pass score of 30, minimum pass percentage 75). For the remaining part of the sequence more relaxed thresholds were applied (minimum phred pass score 10, minimum pass percentage 50). Approximately 2,000,000 sequences per individual were identified and used in marker calling. Finally, identical sequences were collapsed into “fastqcoll” files. The fastqcoll files were used in the secondary proprietary pipeline (DArTsoft14) for SNP and *in silico* DArT (presence/absence of restriction fragments in the representation; PA loci) calling. DArTsoft14 implements a "reference-free" algorithm. All unique sequences from the set of FASTQCOL files are identified, and clustered by sequence similarity at a distance threshold of 3 base variations using an optimised (fast) clustering algorithm (in many cases over 1 billion sequences are clustered within minutes). The sequence clusters are then parsed into SNP and *in silico* DArT markers utilising a range of metadata parameters derived from the quantity and distribution of each sequence across all samples in the analysis. Additionally, a high level of technical replication is included in the DArTseq genotyping process, which enables reproducibility scores to be calculated for each candidate marker. The candidate markers output by DArTsoft14 are further filtered on the basis of the reproducibility values, average count for each sequence (sequencing depth), the balance of average counts for each SNP allele, and the call-rate (proportion of samples for which the marker is scored).

### Sex-linked loci selection

We assessed sex-linkage for both dominant (presence / absence of restriction fragments) and co-dominant (single nucleotide polymorphism, SNP) markers. Each population was analysed separately. Genotypes from the presence / absence (PA) dataset consist of either ‘0’, ‘1’ or ‘-‘, representing fragment absence, presence or putative heterozygosity, respectively. Genotypes from the SNP dataset consist of either ‘0’,’1’,’2’ or ’-‘ representing genotypes homozygous for the reference allele (the most common allele), homozygous for the SNP allele, heterozygous, and homozygous for a null allele (absence of the fragment in the genomic representation), respectively. To evaluate sex linkage, homogeneity of genotypes for all loci between males and females within each population was assessed by Fisher’s exact test using ‘fisher.test’ in R (R Development Core Team 2017) from the ‘stats’ package. *P* values were corrected for false discovery rate by Benjamini and Yekutieli method (Benjamini and Yekutieli, 2001). We assessed sex-linkage amongst the SNP loci under two models. The null exclusive model was conducted with SNP homozygous null genotypes removed. Under this model, sex-linked genotypes that present as homozygous in one sex and heterozygous in the other are expected. Subsequently, we conducted a null inclusive model with SNP homozygous null genotypes_included. Under this model, additional sex-linked genotypes that present as null in one sex and homozygous in the other are expected. The genotypes of all individuals for the sex-linked loci were examined for association with XY and ZW heterogamety. Specifically, XY heterogamety is characterised by PA loci with restriction fragments present in males and absent in females. SNP loci present as homozygous in females (for either the reference or SNP allele) and heterozygous in males under the null exclusive model, or homozygous null in females and homozygous for either allele in males under the null inclusive model. The reciprocal is true for ZW heterogamety. We calculated the probability of loci randomly associating with sex as per Lambert, et al. (2016) to ensure sample sizes were adequate. The PA and SNP markers fitting the null exclusive model were assessed on their ability to discriminate between the sexes of both populations using a Hamming distance matrix calculated using a custom R script with null genotypes removed.

### Comparative linkage disequilibrium (LD) analysis

We used linkage disequilibrium network analysis on a subset of sex-linked SNP loci to examine linkage disequilibrium (LD) within the two populations. LD between two loci occurs when recombination is suppressed along the length of DNA that separates them, and is a hallmark of sex chromosome development (Marshall-Graves 2006). Thus, the number and identity of SNPs in LD with sex-linked SNPs in each population will provide a comparative representation of the sex-determining regions in each population. For this analysis, only SNPs polymorphic in both populations (minor allele frequency > 0.05) were considered. A different but perfectly sex-linked SNP locus (all females homozygous and all males heterozygous) was chosen from each population, along with 100 randomly selected non-sex-linked loci. LD between each of these SNPs and all other SNPs in the dataset was calculated using Genepop V4 (Rousset 2008). SNPs in significant LD (Benjamini and Yekutieli adjusted p value <0.05) were taken for linkage disequilibrium network analysis within their respective population (highland n = 576, lowland n = 618) using the genetics (Warnes, et al. 2013) and LDna (Kemppainen, et al. 2015) packages in R. Parameters for cluster emergence were |E| (the minimum edges or number of connections between loci) set at 20 and phi (factor used to determine the minimum observed change in R^2^ allowed when adding new loci to a cluster) set at 2. Resulting clusters were plotted using the igraph package in R (Csardi, et al. 2006).

## Results

### Sex-linked loci

After DArTsoft14 filtering, DArTseq returned 20,813 presence / absence (PA) loci and 32,663 SNP loci for *Niveoscincus ocellatus.* After correction for false discovery, Fisher’s exact test revealed loci with a non-homogeneous distribution of genotypes between the sexes common to both populations; 152 PA and 54 SNP (p <0.001 to 0.003; supplementary tables S1, S2 and S3). For our sample size of 88 individuals, the probability of any single locus randomly associating with sex is 1.7 × 10^−22^ and therefore it is highly unlikely that any of our loci are sex linked by chance. Of the 152 PA loci, three are perfectly sex-linked across both populations with the remainder having less than 16% of individuals with genotypes deviating from perfect sex-linkage.

Of the 54 sex-linked SNP loci, 21 (supplementary table S2) emerged from the null exclusive model and are homozygous in females and heterozygous in males. Seven of these SNPs are perfectly sex-linked across both populations with the remainder sex-linked in at least 77% of individuals. The remaining 33 (supplementary table S3) emerged from the null inclusive model. These loci present more like presence / absence loci because the majority of females possess a homozygous null genotype and the majority of males are homozygous for either the SNP or reference allele. One of these loci is perfectly sex-linked across both populations with the remainder having less than 17% of individuals with genotypes deviating from perfect sex-linkage.

Fisher’s exact test also revealed PA and SNP loci that are sex-linked in one population only. Sixteen PA, 12 SNP loci from the null exclusive model and eight SNP loci from the null inclusive model in the highland, and 20 PA, 16 SNP loci from the null exclusive model and five from the null inclusive model in the lowland population, exhibited non-homogeneity of genotypes between males and females uniquely within their respective populations (p<0.001 to 0.012; supplementary tables S1, S2 and S3). Proportional pairwise Hamming’s distances between males and females (Figure 1) using the population-specific PA and SNP loci (null exclusive model), demonstrate that they reliably reveal an organism’s phenotypic sex within that population only. Highland males and females are on average 89.7% and 92.8% dissimilar from one another (Highland SNP and PA loci, respectively). Lowland males and females are on average 89.4% and 89.2% dissimilar from one another (Lowland SNP and PA loci, respectively).

**Figure 1.**
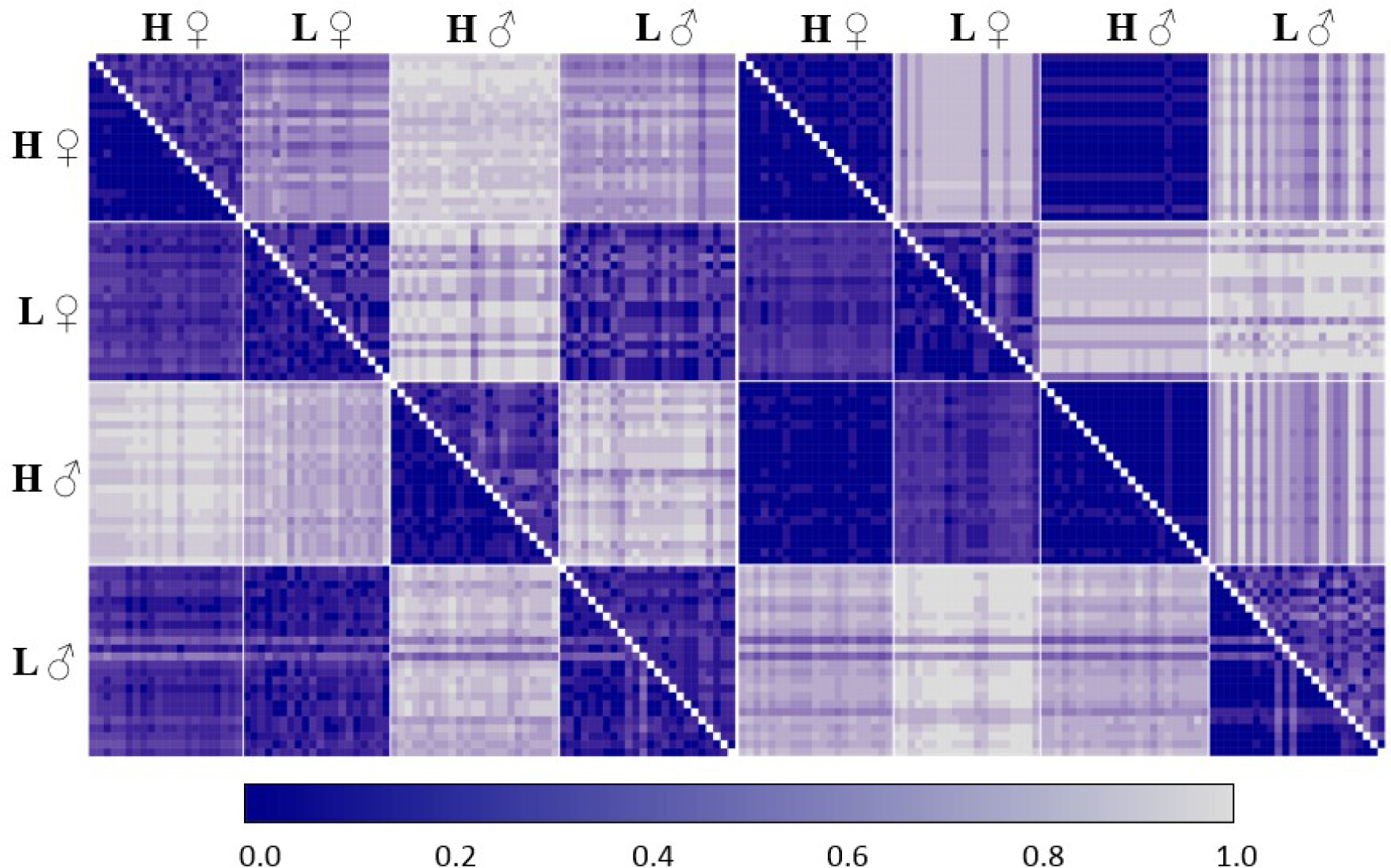
Hamming’s proportional distance among Niveoscincus ocellatus individuals of highland (H) and lowland (L) populations for sex-linked loci unique to the highland (left panel) and lowland (right panel) populations. Presence Absence (PA; lower segment) and SNP (upper segment). Highland PA n=16, lowland PA n=20, highland SNPs n=12, lowland SNPs n=16.

All sex-linked SNP loci specific to a given population (hereafter the “source population”) were also genotyped in the other population (hereafter the “reciprocal population”). Genotypes in the reciprocal population were not sex-linked because an allele was either absent in both males and females (all individuals monoallelic) or both alleles were homogenously distributed among the sexes. Where an allele was absent from one sex it was always missing from the females of the reciprocal population (homozygous null), with males possessing one of the source population alleles. These loci were all recovered as sex-linked in the reciprocal population under the null inclusive model. For the population-specific sex-linked PA loci, in the reciprocal population the restriction fragment in question was either absent in all individuals, present in all individuals, or present at homogeneous frequencies between the sexes. Males and females are more dissimilar in the lowland (21.8% and 16.3%, SNP and PA loci respectively) than highland (2.6% and 4.4%) population based on sex-linked loci from the reciprocal population (Figure 1).

Sex-linked genotypes assort in a manner consistent with XY heterogamety: PA loci present in males, absent in females; SNPs are either heterozygous in males and homozygous in females or males are monomorphic and females are homozygous null, consistent with Y chromosome specific sequence. Exceptions are five loci in the lowland population. One lowland PA locus is absent from all males and 45% of females (homogeneity of genotypes, p = 0.002). Four SNP loci are homozygous for every male individual, but for both alleles at each locus, while most females are heterozygous, but with homozygotes also observed for both alleles at each locus (homogeneity of genotypes p<0.001). These five loci were recovered in the highland population, but the genotypes of the PA locus are homogeneous between the sexes (p>0.05) and those of the SNPs are homozygous for the reference allele in all individuals.

### Comparative LD analysis

Linkage disequilibrium network analysis (LDNa) resolved a sex-linked cluster consisting of 32 SNP loci connected via 411 edges in the highland population (12.8 edges per locus; Figure 2). 175 non-sex-linked loci connected by 213 edges described a non-sex-linked cluster in this population (1.2 edges per locus). In the lowland population, LDNa resolved a sex-linked cluster with 34 SNP loci connected via 235 edges (6.9 edges per locus; Figure 2) and a non-sex-linked cluster containing 17 loci connected via 22 edges (1.3 edges per locus). The 21 common sex-linked SNP loci (Figure 2 - **a** to **u**; supplementary table S2) appear in both the highland and lowland sex-linked clusters but vary in the degree to which they associate with the perfectly sex-linked locus for that population and each other. The non-sex-linked clusters from each population have no loci in common.

**Figure 2.**
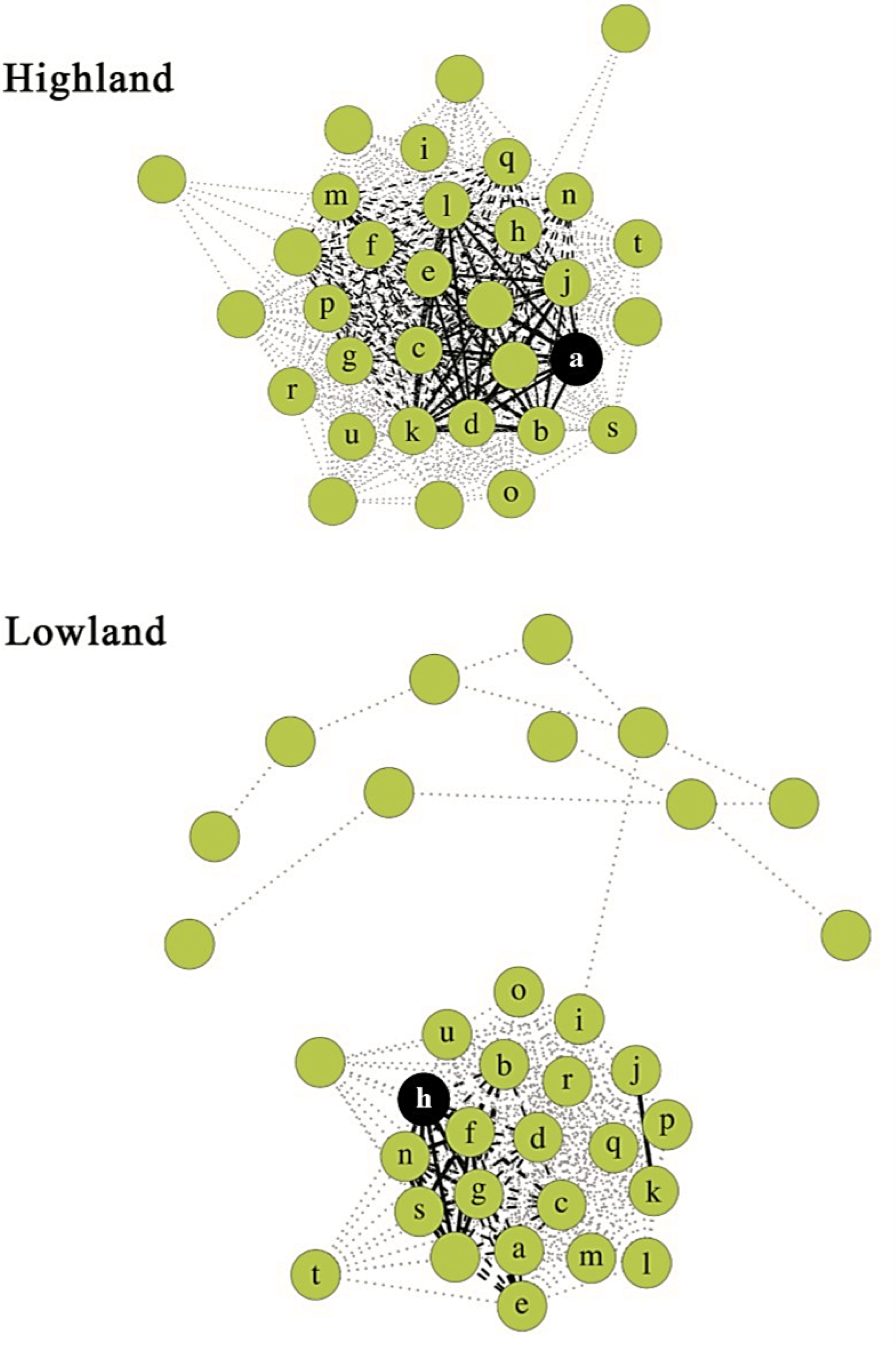
Linkage disequilibrium network plot of sex-linked clusters from the highland and lowland populations of Niveoscincus ocellatus. Green circles indicate sex-linked SNPs (n = 32 in the highland, n = 34 in the lowland populations); ‘a’ to ‘u’ denote 21 loci sex-linked in both the highland and lowland population. The perfectly sex-linked locus for each cluster is in black. Black solid edges have an R2 > 0.99, black dashed 0.99>R2>0.80, grey dotted R2<0.80.

## Discussion

The subtle molecular differences in sex-linked markers between highland and lowland populations indicate that few changes are required for transitions between sex determination modes, and is also compatible with the short timeframe (<1 Myr) across which these populations have diverged (Cliff, et al. 2015). We identified a similar number of loci associated with phenotypic sex in both populations of *N. ocellatus,* despite their different mechanisms of sex determination. This was surprising because lowland offspring sex ratios are correlated with temperature (Wapstra, et al. 2004), and models predict the loss of genes surrounding sex determination in this population, or the gain of such genes in the highland population (Pen, et al. 2010). In addition to a conserved set of sex-linked loci, population-specific sex-linked loci are also present in each population, highlighting the importance of a genetic contribution to sex in the lowland as well as the highland. Sex ratios in the lowland are 50:50 across a pivotal temperature range (Wapstra, et al. 2009), likely facilitated by random assortment of sex determining genes at meiosis and maintained due to frequency-dependent selection (Fisher 1958). Developmental temperatures outside this pivotal range provide sex-specific fitness advantages (Pen, et al. 2010), and there has been selection for a temperature mediated dosage component to sex determination both above and below the pivotal temperature in this population. *N. ocellatus* is an XY GSD species with the lowland population possessing a temperature over-ride in sex determination (GSD+EE); the maintenance of this mixed system in the lowland likely representing an adaptive optimum.

The presence of sex-linked markers in a population exhibiting TSD can be explained by mechanisms that only occasionally over-ride GSD, and this is consistent with the low but significantly temperature-related deviations in sex ratio observed in the lowland population (Cunningham, et al. 2017; Wapstra, et al. 2004). Both differential mortality and differential fertilization via cryptic female choice have been implicated in other taxa (Burger, et al. 1988; Eiby, et al. 2008; Olsson, et al. 2011), but have been ruled out in this species (Wapstra, et al. 2004). This leaves sex reversal as the most likely explanation for the sex ratio biases observed. Sex reversals can occur via temperature sensitive gene dosage and in reptiles usually occurs in the homogametic sex (Ezaz, et al. 2009b; Holleley, et al. 2015; Quinn, et al. 2007; Quinn, et al. 2011). Explanations for this centre around a gene or gene product present on the homogametic chromosome and therefore present as one copy in one sex and two in the other. Temperature-sensitive, dosage-dependent expression of this gene or activity of its product can result in the homogametic genotype not reaching the threshold for sexual phenotype and becoming sex reversed. Male biased sex ratios in *N. ocellatus*, as observed in colder conditions, fit this pattern if sex determination in this species occurs via a feminizing gene on the X chromosome with sex reversed males (XX genotype) resulting from temperature sensitivity of this gene. When gene product fails to reach the required threshold to produce a female, a male is instead produced from this genotype. Female biased sex ratios, as observed in warmer conditions, could result from the over-expression of this feminizing allele in the XY genotype. Sex reversal in the heterogametic sex is thought to be unfavourable when sex chromosomes are highly heteromorphic; mating between sex reversed XY females and XY males producing YY progeny - unviable if there are necessary developmental genes on the X chromosome (Quinn, et al. 2011). However, sex reversal of the XY genotype to female in systems with homomorphic sex chromosomes is theoretically possible (Sarre, et al. 2004) and could explain the observed differences in recombination suppression in the two populations of *N. ocellatus.*

The ratio of female to male recombination rate varies considerably across taxa (Berset-Brändli, et al. 2008; Coimbra, et al. 2003; Perrin 2009) even in taxa without sex chromosomes (Isberg, et al. 2006), and is a function of phenotypic rather than genetic sex. Recombination between sex chromosomes can therefore occur in individuals that have been sex reversed (Perrin 2009). In an XY system the X and Y chromosome can undergo recombination at meiosis in sex reversed XY females, resulting in reduced associations between alleles on the Y. This interrupts the progressive degeneration of the Y chromosome because recombination suppression is necessary to keep alleles beneficial for one sex together. Sex reversals are described in reptiles, amphibians and fish (Alho, et al. 2010; Holleley, et al. 2015; Shao, et al. 2014), but have yet to be described in *N. ocellatus.* The occurrence of sex reversed females in the lowland population would explain lower linkage disequilibrium between the sex-linked SNP loci in this population.

Although the number of sex-linked SNPs and PA loci is similar in both populations, the presence of population-specific sex-linked variation nevertheless supports population divergence in the molecular mechanism surrounding sex determination. The degree of recombination suppression occurring amongst the 21 shared sex-linked markers also differs among populations, indicating sex chromosomes at different developmental stages. Sex chromosomes in the highland population are likely more differentiated than those in the lowland because of the lower independence of genotypes between sex-linked loci in this population. This lower independence manifests as both higher LD between loci and a greater number of connections among the 21 shared loci, suggesting a region that is tightly bound and likely travelling as a complete unit during meiosis because of suppressed recombination. Many taxa (e.g., Ratite birds and Boid snakes) maintain recombination along much of the length of their sex chromosomes (Vicoso, et al. 2013a; Vicoso, et al. 2013b). Recombining sex chromosomes are advantageous as deleterious alleles are purged from the Y (or W) chromosome (Bachtrog, et al. 2014; van Doorn, et al. 2010). Recombination between the X and Y may contribute to the maintenance of a mixed system in the lowland population where temperature and genetics interact to determine sex (Sarre, et al. 2004) via the presence of sex reversed females.

Here we describe sex-linked genetic sequence in *Niveoscincus ocellatus.* The majority of sex-linked markers observed in this study were shared between populations, indicating inheritance from a common ancestor; those not shared may indicate independent gain or loss in a population. A thorough examination of sex determination across this genus using these loci will reveal the ancestral state of sex determination in *N. ocellatus* and whether population divergence in sex determination occurs elsewhere in the genus. Further, these loci can be used to assess the role of sex reversal in the transition in sex determination mode in this species, for cytological examination of the karyotype of this genus and to uncover the sex determining locus in Scincidae. Screening our archival samples, collected over more than a decade, with these sex-linked markers will be invaluable in capturing the tempo and mechanism of evolutionary transitions between modes of sex determination in reptiles.

## Acknowledgements

We thank the following people: G. Cunningham and L. Fitzpatrick for field collection and valuable contributions and discussions, A. Kilian and Diversity Arrays Technology for sequencing support. We thank the Holsworth Wildlife Research Endowment and the Australasian and Pacific Science Foundation for their contributions to this research. All guidelines and procedures for the use of animals were approved by the University of Tasmania animal ethics committee (no. A0012087). This work was supported by the Australian Research Council (FT110100597).

